# Stabilization of H5 highly pathogenic avian influenza hemagglutinin improves vaccine-elicited neutralizing antibody responses

**DOI:** 10.1101/2025.07.30.667762

**Authors:** Annie Dosey, Bernadeta Dadonaite, Rebecca A. Gillespie, Elizabeth M. Leaf, Matthew J. Vukovich, Jackson McGowan, Emily Grey, Hiromi Muramatsu, Rachel H. J. Jun, Norbert Pardi, Masaru Kanekiyo, Jesse D. Bloom, Neil P. King

## Abstract

Transmission of highly pathogenic avian influenza from H5 clade 2.3.4.4b has expanded in recent years to infect large populations of birds and mammals, heightening the risk of a human pandemic. Influenza viruses adapted to transmission in birds and some other animals tend to have a less stable hemagglutinin (HA) than seasonal influenza viruses, enabling membrane fusion at comparatively high pH levels. Here, we combine five mutations within H5 HA that dramatically increase its melting temperature and promote stable closure of the HA trimer. Structural analysis by cryo-electron microscopy revealed that the stabilizing mutations create several new hydrophobic interactions, while maintaining local HA structure. We found that vaccinating mice with stabilized H5 HA immunogens resulted in higher hemagglutination inhibition and neutralization titers than non-stabilized comparators. Epitope mapping of vaccine-elicited polyclonal antibody responses using negative stain electron microscopy and deep mutational scanning showed that site E on the side of the HA receptor binding domain was immunodominant across all groups; however, the stabilized immunogens shifted responses toward the receptor binding site (RBS), eliciting a higher proportion of neutralizing antibodies. These findings highlight that H5 HA-stabilizing mutations enhance the quality of antibody responses across different vaccine formats, underscoring their potential to improve pandemic preparedness vaccines targeting viruses from this widely circulating clade.

**One Sentence Summary:** Vaccinating with stabilized H5 HA increases the proportion of humoral responses against neutralizing RBS epitopes.

## INTRODUCTION

Since 1996, highly pathogenic avian influenza (HPAI) H5 viruses have been spreading at a large scale amongst both wild and domesticated birds, with spillover infections occurring in several mammalian species, including humans (*1*). By 2021, clade 2.3.4.4b had emerged as the dominant global strain and was introduced to North America through wild bird migration (*2, 3*). In 2024, large H5 outbreaks in both dairy cattle and domesticated poultry across the United States devastated both industries and led to several dozen infections in humans (*4–7*). Large-scale infections of HPAI H5 have been observed in several other mammals as well, including sea lions, foxes, llamas, and cats, underscoring its adaptability (*8*). Underlying these outbreaks, recent studies suggest that H5 transmission continues to increase through multiple evolutionary mechanisms (*9, 10*). This is especially concerning given *in vitro* evidence that only a single mutation in an avian H5 HA is needed to bind to a human receptor (*11, 12*). The persistent evolution and broad circulation of HPAI H5 across multiple animal species pose an ongoing threat of a potential human pandemic.

Given the high evolutionary rate of HPAI H5, a quick turnaround time to produce new vaccines is necessary. Greatly improved production speed has been achieved through the advent of mRNA vaccines, which enable a single manufacturing process for any sequence (*13, 14*). Already, several studies employing membrane-anchored H5 HA mRNA vaccines have shown elicitation of neutralizing antibodies in mice and ferrets, and human clinical trials have been initiated (*15–20*). Recent work from our lab combining mRNA vaccines with the enhanced immunogenicity of multivalent antigen display showed that an mRNA-launched computationally designed protein nanoparticle displaying 60 copies of the SARS-CoV-2 spike receptor binding domain (RBD) elicited significantly higher neutralization titers in mice compared to membrane-anchored spike (*21*). These findings highlight the potential of integrating mRNA technology with protein nanoparticle display to swiftly generate effective vaccines against rapidly evolving viruses like HPAI H5.

Despite the continued focus on HA as the primary influenza vaccine antigen—and unlike many other class I viral fusion proteins(*22*)—there has been little work examining if increasing HA protein stability improves immunogenicity. Only recently, Milder et al. showed that the poor expression and monomeric nature of several HA subtypes when expressed without trimerization domains, including H5, can be improved through structure-based vaccine design (*23*). Even with the addition of foldon or the trimeric nanoparticle component I53_dn5B as trimerization domains, considerable instability and breathing of the HA was observed. To improve HA stability, they introduced hydrophobic mutations in two pH-switch regions conserved across all major influenza A subtypes that dramatically improved expression and trimerization. In a separate study, stability-enhancing mutations in H5 HA were identified through successive heat treatments of a virus library incorporating random mutations (*24*). Further evidence of the marked instability of H5 HA is seen in the increased pH of fusion of this subtype as compared to seasonal influenza strains (*25*). Perhaps relatedly, an inactivated H5 vaccine has been observed to be dramatically less immunogenic as compared to the seasonal inactivated vaccine in ferrets (*26*). Despite this evidence for H5 protein instability, as well as the previous identification of stabilizing mutations, no work assessing the impact of stabilization on immunogenicity *in vivo* has been reported.

Here, we introduced mutations into the clade 2.3.4.4b influenza H5 HA protein to improve its stability and compared the wild-type and stabilized HA in several protein and mRNA vaccine formats. We found that delivering stabilized protein nanoparticle and membrane-anchored H5 HAs as mRNA-LNPs elicited a higher proportion of RBS-directed responses and the most potent hemagglutination inhibition (HAI) and neutralizing activity.

## RESULTS

### Stabilizing mutations increase melting temperature and trimeric closure of H5 HA

To stabilize the prefusion conformation of HPAI A/Texas/37/2024 H5 HA, combinations of previously identified mutations from multiple studies were tested for increased thermal stability using differential scanning fluorimetry (nanoDSF) (*11, 23, 24, 27*). The H5 HA variant with the highest melting temperature (T_m_) incorporated the following mutations within the HA2 domain: H355F/K380M/R397M/L418I/E432L (based on mature H3 HA numbering(*28*)) (Fig. 1A). Hereafter we refer to this stabilized HA as H5-FMLMI, as it contains mutations to these amino acid identities at stabilized sites. This H5 HA was genetically fused via flexible (GS)_n_ linkers to a foldon trimerization domain or the previously described one-component *de novo* nanoparticle RC_I_1 (Fig. 1B and table S1) (*29*). Additionally, all H5 HAs used in this study contain the Y98F mutation(*30, 31*) and have a mutated polybasic cleavage site, resulting in HA0 constructs. The T_m_ of H5-FMLMI-foldon was 15-20°C higher than the wild-type H5-foldon at both pH 8.0 and pH 5.5 (Fig. 1C). The T_m_ of H5-FMLMI-RC_I_1 was 28°C and 40°C higher than H5-RC_I_1 at pH 8.0 and pH 5.5, respectively. The low T_m_ of 35°C for H5-RC_I_1 at pH 5.5, compared to 59°C for H5-foldon, indicated that this scaffold is relatively unstable at low pH. However, this instability was mitigated by the stabilizing mutations within the HA antigen, which resulted in similar T_m_ values at pH 5.5 for H5-FMLMI-foldon and H5-FMLMI-RC_I_1.

**Fig. 1.**
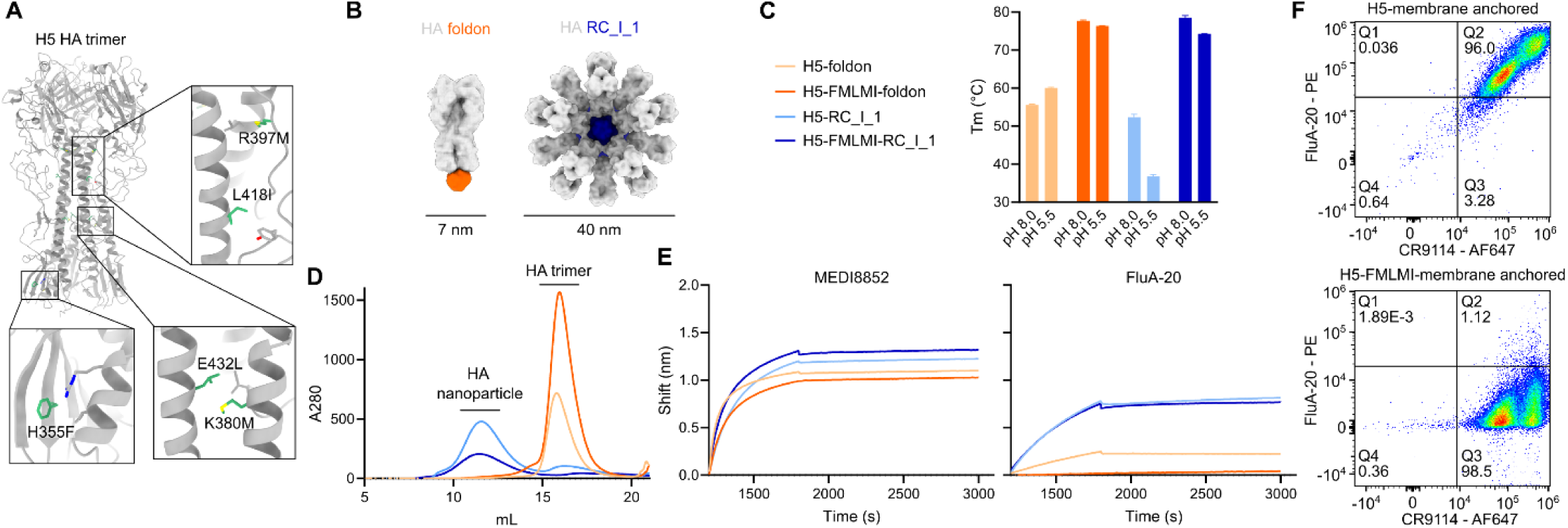
Stabilizing mutations within H5 increase thermostability and reduce binding of internally-directed mAb. (A) Stabilizing mutations H355F/K380M/R397M/L418I/E432L (green) highlighted within the A/Texas/37/2024 crystal structure (PDB ID: 9DIQ). Key interacting side chains are shown in gray. (B) Surface models of HA foldon trimer and HA RC_I_1 one-component nanoparticle. (C) Melting temperatures of H5 HA constructs at pH 8.0 and pH 5.5 as measured by nanoDSF. (D) SEC chromatograms of H5 HA constructs. Same legend as panel (C). (E) BLI of H5 HA constructs against anti-stem mAb MEDI8852 and anti-trimer interface mAb FluA-20. Same legend as panel (C). (F) Flow cytometry plots of CR9114 and FluA-20 binding to membrane-anchored H5 HA constructs expressed on the surface of HEK293F cells.

Final stabilized and unstabilized H5-foldon and -RC_I_1 nanoparticle constructs were scaled up for production in HEK293F cells. Following immobilized metal affinity chromatography (IMAC), proteins were further purified by SEC, resulting in chromatograms with single elution peaks at expected retention volumes for all four constructs (Fig. 1D). Final protein yields per liter of cell culture were 12.8 mg/L for HA-foldon, 26.8 mg/L for H5-FMLMI-foldon, 10.4 mg/L for H5-RC_I_1, and 4.4 mg/L for H5-FMLMI-RC_I_1. SDS-PAGE of these elution peaks showed a single band that migrated at the expected molecular weight for all constructs (fig. S1A).

Negative-stain electron microscopy (nsEM) micrographs and 2D class averages showed both H5 foldon constructs formed trimeric HAs and both H5 RC_I_1 constructs formed intact, monodisperse nanoparticles (fig. S1B). Dynamic light scattering (DLS) showed the expected size for both H5 foldon constructs, but both H5 RC_I_1 nanoparticles had a hydrodynamic radius of 30 nm, 10 nm smaller than the H5 RC_I_1 design model (fig. S1C) (*29*). The mass calculated using mass photometry (MP) of H5-FMLMI-RC_I_1 was consistent with approximately 8 trimers per nanoparticle (fig. S1D). The nsEM, DLS, and MP together suggest that both H5 RC_I_1 constructs formed octahedra rather than the intended icosahedra.

Bio-layer interferometry (BLI) using two anti-RBS mAbs (310-12D03 and 310-1H02), three lateral patch mAbs (310-7D11, 326-289.74, and 326-366.26), and the anti-stem mAb MEDI8852 (*32*) all showed similarly high binding for all four H5 HA constructs (Fig. 1E and fig. S1E). FluA-20 is a mAb that binds to the interface between HA monomers within the trimer, so it is only able to bind to partially open HA trimers (*33*). Similar binding was observed for FluA-20 against both H5-RC_I_1 and H5-FMLMI-RC_I_1, suggesting incomplete trimer interface closure in this H5 nanoparticle despite the addition of the stabilizing mutations (Fig. 1E). We note that partial, but incomplete, trimer closure may not be detected using this experimental approach, since the presence of even a rare single open HA trimer per nanoparticle would result in binding of the entire particle. By contrast, FluA-20 bound to H5-foldon but not H5-FMLMI-foldon, suggesting that these stabilizing mutations induce stable closure of the trimer interface in this construct. Conformation-specific mAbs were also used to probe the antigenicity of membrane-anchored H5 HA on the surface of HEK293F cells (Fig. 1F). High anti-stem mAb CR9114 (*34*) and FluA-20 binding was observed for H5-membrane-anchored, while high CR9114 and minimal FluA-20 binding was observed for H5-FMLMI-membrane-anchored. These results indicate that these H5 membrane-anchored constructs express similarly, but that the head domains of the wild-type H5 HA are open or monomeric while the FMLMI stabilizing mutations encourage closed trimer formation.

### Cryo-electron microscopy (cryo-EM) reconstruction of H5-FMLMI-foldon

To understand how the FMLMI mutations contributed to H5 HA stabilization and its conformation, we determined a 3.2-Å resolution reconstruction of H5-FMLMI-foldon by cryo-electron microscopy (Fig. 2A, fig. S2, A and B, and table S2). The backbone RMSD to a TX24 H5 HA crystal structure (PBD ID: 9DWE) was 3.5 Å over the entire trimer, due to the H5-FMLMI-foldon HA1 and HA2 domains shifting closer to one another, resulting in a more compact structure overall (Fig. 2B). Despite this relatively high RMSD over the entire trimer, the RMSD was lower when aligning on individual epitopes: 1 Å RMSD for the stem epitope and 0.6 Å RMSD for the RBS epitope (Fig. 2B). This is consistent with the similar binding we observed between stabilized and unstabilized H5 HA constructs *in vitro* for anti-RBS and anti-stem mAbs (Fig. 1E and fig. S1E). We conclude that while the stabilizing mutations induce an overall rigid body shift between domains, the local structure of HA remains preserved. Further analysis of the stabilizing mutations in the cryo-EM reconstruction showed formation of several hydrophobic interactions (Fig 2, C and D). R397M replaces polar interactions of R397 with N410 and N408 with hydrophobic contacts to L409. L418I encourages hydrophobic packing with Y308. E342L and K380M replaced this internal salt bridge with hydrophobic interactions to L428, I384, and M30. Finally, H355F replaced a pH-dependent ionic interaction of H355 to R484 with a hydrophobic contact to M478. Similar substitutions that create new hydrophobic interactions have been previously observed in other HA stabilization efforts(*23*).

**Fig. 2.**
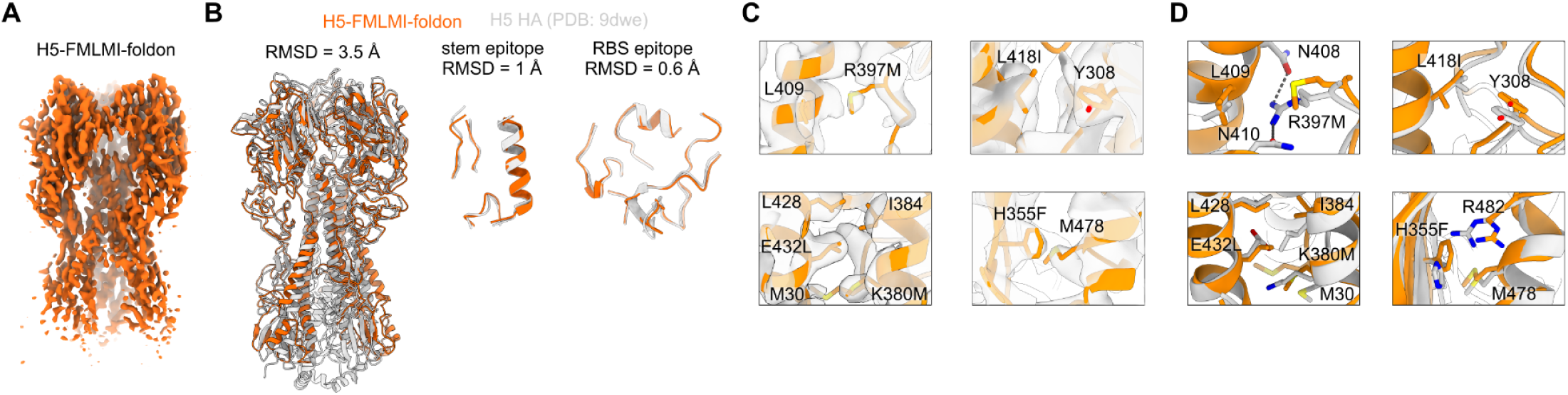
Cryo-EM of H5-FMLMI-foldon. (A) Cryo-EM reconstruction of H5-FMLMI-foldon. (B) Superimposition of the cryo-EM model of H5-FMLMI-foldon and the A/Texas/37/2024 crystal structure (PDB ID: 9DWE) for the H5 HA ectodomain (left), the central stem epitope (center), and the RBS epitope (right). (C) Close-ups of density for the FMLMI mutations in the H5-FMLMI-foldon reconstruction with the built model. (D) Close-ups of the FMLMI mutations and interacting residues in the cryo-EM model of H5-FMLMI-foldon (orange) superimposed with the A/Texas/37/2024 crystal structure (PDB ID: 9DWE; gray).

### H5 HA-stabilizing mutations increase neutralizing antibody responses in mice

Next, we evaluated these stabilized H5 HA constructs as both protein- and nucleoside-modified mRNA-LNP-delivered vaccines in BALB/c mice. Mice were subcutaneously immunized at weeks 0 and 4 with 21.8 pmol total protein (1.37 μg for H5 foldon constructs and 1.5 μg for H5 nanoparticles) formulated with AddaVax or 1 μg mRNA-LNP (Fig. 3A). Vaccine-elicited serum antibody responses were assessed two weeks post-prime and post-boost.

**Fig. 3.**
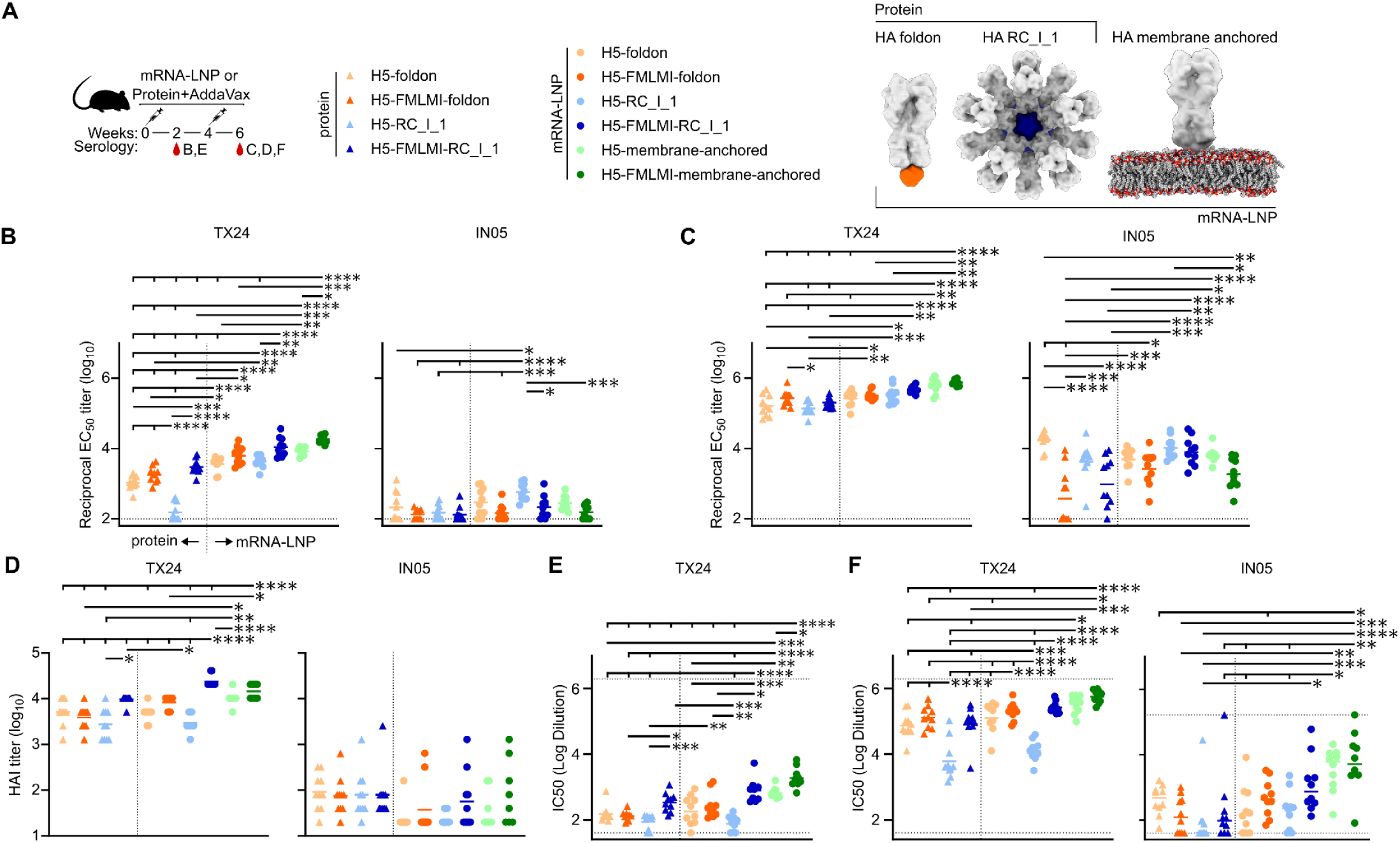
Vaccine-elicited Antibody Responses in Mice Immunized with H5 protein- and mRNA-LNP-delivered constructs (A) H5 mouse immunization schedule and groups. Molecular models of immunogens are not to scale. (B-C) ELISA binding titers against vaccine-matched TX24 and vaccine-mismatched IN05 in immune sera at B. week 2 and C. week 6. (D) HAI titers in immune sera at week 6. (E-F) Microneutralization titers in immune sera at E. week 2 and F. week 6. Each symbol represents an individual animal, and the geometric mean of each group is indicated by the bar (N = 9-10 mice/group). Statistical significance was determined using one-way ANOVA with Tukey’s multiple comparisons test; *, p < 0.05; **, p < 0.01; ***, p < 0.001; ****, p < 0.0001.

To assess serum binding, vaccine-matched TX24 and vaccine-mismatched A/Indonesia/175H/2005 (IN05) HAs genetically fused to the trimeric nanoparticle component I53_dn5B (*35, 36*) were used in ELISAs. Both post-prime and post-boost, there were slightly higher TX24 binding titers in all stabilized groups compared to the similar construct without stabilization (Fig. 3, B and C). Overall, the unstabilized H5 nanoparticle had lower TX24 binding, HAI, and neutralizing activity than the other groups (Fig. 3, B to F). By contrast, the mRNA-LNP-delivered stabilized nanoparticle elicited significantly higher TX24 HAI titers than all other groups (Fig. 3D). The highest TX24 neutralizing activity, both post-prime and post-boost, was seen in the mRNA-LNP-delivered stabilized nanoparticle and both membrane-anchored groups, with the stabilized membrane-anchored group showing significantly higher post-prime neutralization than the unstabilized group (Fig. 3, E and F). Compared to TX24, the opposite trend in binding titers was observed for IN05, where the unstabilized groups all had higher binding titers than their stabilized counterparts (Fig. 3, B and C). By contrast, IN05 HAI and neutralization titers were comparable or trended slightly higher in stabilized constructs compared to their unstabilized counterparts (Fig. 3, D to F). The lower binding but higher neutralization IN05 titers observed in the stabilized groups compared to the unstabilized groups indicates that the stabilized immunogens elicited a greater proportion of functional, vaccine-mismatched antibodies, perhaps by reducing the proportion of trimer interface-directed antibodies. Taken together, these results demonstrate that the stabilizing mutations consistently increased the elicitation of functional antibody responses across all constructs, although not all comparisons were statistically significant.

With stabilizing mutations, the membrane-anchored and nanoparticle groups were observed to have as good or significantly better HAI and neutralizing activity than the H5-foldon trimer in all instances (Fig. 3, D and F). Slightly lower anti-stem titers against stabilized-stem TX24 (ssTX24) were elicited in the membrane-anchored groups as compared to the nanoparticle groups, though the opposite trend was observed against the TX24 HA ectodomain, suggesting that a greater proportion of the antibodies elicited by the nanoparticle groups are against the HA stem (Fig. 3C and fig. S3B). However, the greatest proportion of cross-neutralizing antibodies was observed in the stabilized membrane-anchored group, since it had the lowest IN05 binding titers out of all mRNA-LNP groups and the highest IN05 neutralization. Thus, the increased avidity provided by the nanoparticle and membrane-anchored constructs enhanced immune responses as compared to foldon trimers, though the nature of these improvements differed between formats.

mRNA-LNP-delivered constructs overall elicited comparable or improved binding, HAI, and neutralization titers as compared to the same construct delivered as protein. All mRNA-LNP-delivered constructs elicited significantly higher post-prime TX24 binding titers than their protein-delivered counterparts (Fig. 3B).

Additionally, mRNA-LNP-delivered H5-FLMI-RC_I_1 elicited significantly higher HAI than protein-delivered (Fig. 3D). Higher anti-scaffold responses were also observed upon vaccine delivery with mRNA-LNP as compared to protein for the nanoparticle groups, while scaffold responses between foldon groups were similar across delivery formats (fig. S3B). Since mRNA-LNP delivery of several different HA constructs resulted in at least comparable and sometimes superior immunogenicity as compared to protein delivery, these results underscore the potential of mRNA-LNP-based vaccine platforms.

### H5 HA-stabilizing mutations increase the proportion of neutralizing RBS-directed responses

We first used nsEM polyclonal epitope mapping (ns-EMPEM) to map the epitopes targeted by vaccine-elicited antibody responses. Purification of vaccine-matched TX24 H5 HA-I53_dn5B bound to polyclonal Fabs purified from serum elicited by the six mRNA-LNP immunogens showed less early-eluting, high molecular weight material for all three stabilized H5 HA constructs as compared to their unstabilized counterparts (Fig. 4A).

**Fig. 4.**
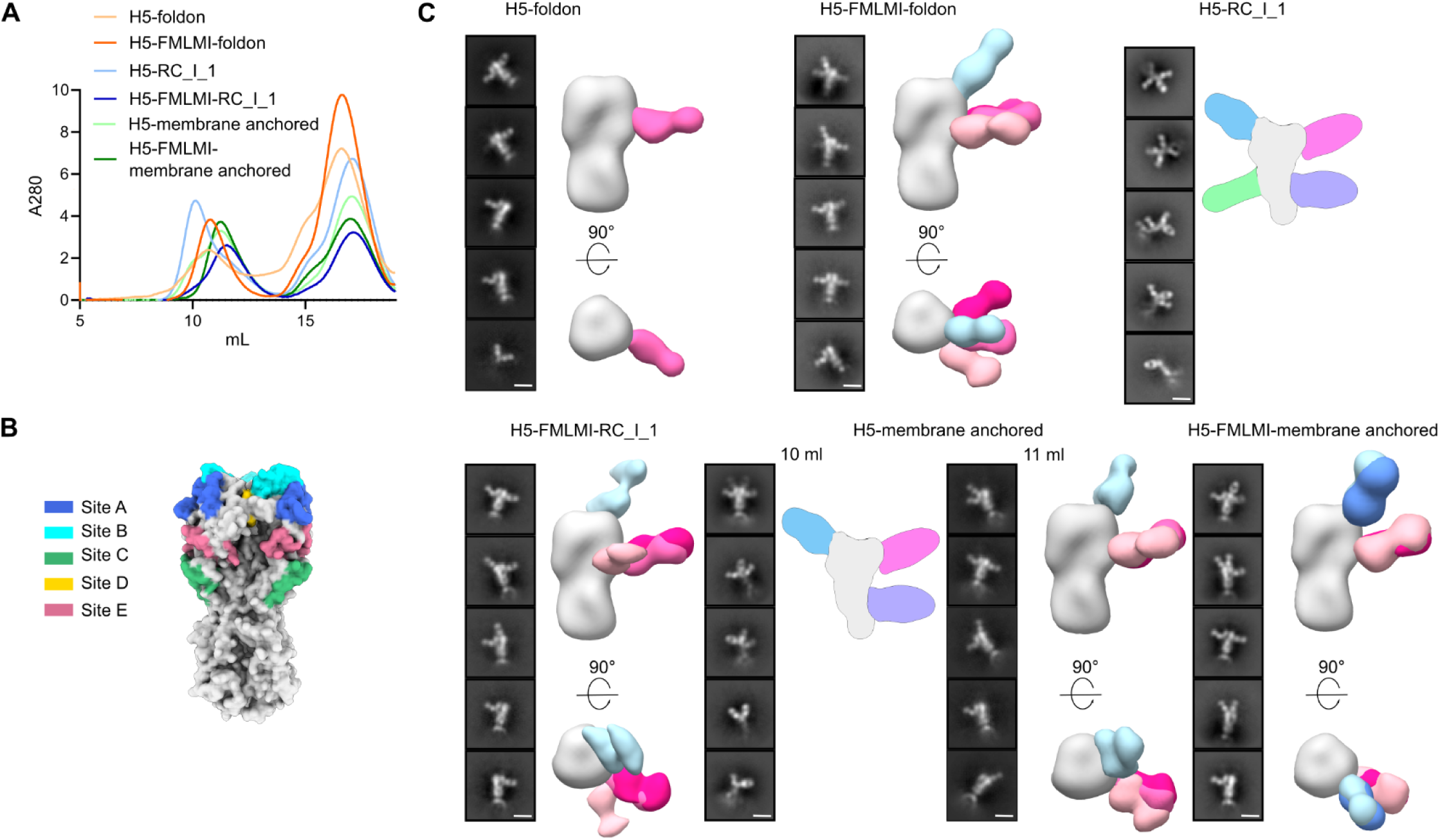
ns-EMPEM of Vaccine-elicited Antibody Responses. (A) SEC chromatograms of TX24 HA complexed with polyclonal Fabs from H5 mouse study at week 6. (B) Locations of conventional H3 HA antigenic sites (*37*). (C) Representative 2D class averages of purified immune complexes in (A). H5-RC_I_1 and the H5-membrane-anchored 10 mL fraction contain a cartoon schematic of a likely 3D model, while all other groups are composite 3D models from ns-EMPEM analysis. Scale bars = 15 nm.

Immune complex peak fractions collected around 10-12 mL were selected for ns-EMPEM analysis. Serum antibodies were observed to bind to previously defined antigenic sites on the HA head (Fig. 4B) (*37*). For the HA-foldon constructs, the vast majority of antibodies elicited by the unstabilized immunogen appeared to target site E, which contains the vestigial esterase domain, while the stabilized immunogen induced responses against both site E and the RBS (Fig. 4C). Additionally, HA-foldon showed a minor population of Fab bound to HA monomer, indicative of responses elicited against internal epitopes that disrupt trimerization, but this was not observed in the HA-FMLMI-foldon group. In the nanoparticle groups, H5-RC_I_1 revealed only Fab-bound HA monomers, with responses against several epitopes along the HA monomer, accounting for the earlier elution of larger complexes in this unstabilized group. By contrast, all Fabs elicited by H5-FMLMI-RC_I_1 were visualized in complex with trimeric HA, targeting both site E and RBS epitopes. Since the unstabilized H5-membrane-anchored immune complex SEC profile showed a large shoulder at 10 mL before a slightly larger peak at 11 mL, ns-EMPEM was performed on both fractions. The majority of complexes in the 10 mL fraction showed multiple Fabs bound to monomeric HA at distinct epitopes, explaining their earlier elution, though a small number of complexes comprising site E- and RBS-directed Fabs bound to trimeric HA were also observed. By contrast, only the latter were observed in the 11 mL fraction. Similarly to the nanoparticle results, stabilization in the H5-FMLMI-membrane-anchored group resulted in no Fab-bound HA monomers and the largest proportion of RBS-directed Fabs out of all groups, in addition to some responses against site E. These data allow us to draw multiple conclusions. First, site E appears to be immunodominant in TX24 HA. Second, stabilization of H5 HA through the introduction of substitutions in HA2 reduces the elicitation of trimer interface-directed antibodies, likely by maintaining closure of the head domain *in vivo* as suggested by the lower FluA-20 binding we observed with these constructs *in vitro*. Finally, stabilization of H5 HA increases the proportion of vaccine-elicited antibodies targeting the RBS.

To compare the epitopes targeted by neutralizing antibodies in the sera from each group of immunized mice, we used a pseudovirus-based deep mutational scanning (DMS) system. Pseudovirus DMS provides a safe and comprehensive method to measure the effects of HA mutations on neutralization by using pseudotyped viral particles that can only undergo a single round of cell entry (*11*). A library of pseudoviruses expressing HA variants covering every possible HA ectodomain amino-acid mutation was incubated with sera from multiple animals from each group that received mRNA-LNP-delivered H5 vaccines, and escape mutations that led to decreased serum neutralization were identified (Fig. 5). For all groups except H5-RC_I_1, escape mapping revealed that regardless of the presence of HA stabilizing mutations, neutralizing antibodies primarily targeted the A and B epitopes at the top of the HA RBD, including the receptor binding site (RBS) (Fig. 5, A and B). The sites with the highest escape for these sera were 129, 145, 158, 169 and 189 (H3 numbering). By contrast, when using sera from animals that received H5-RC_I_1, escape mutations were also observed in site E, at positions 57, 78, 90, and 276 (Fig. 5, A and B). This result is interesting because responses against site E were also clearly visualized in sera from the H5-FMLMI-RC_I_1 group (Fig. 4C). We conclude that lower levels of potently neutralizing RBS-directed antibodies in the H5-RC_I_1 sera revealed the existence of more weakly neutralizing site E-directed antibodies in the DMS assay. Although ns-EMPEM data also showed that HA stabilization also led to a greater proportion of RBS-targeting antibodies for the membrane-anchored and HA-foldon constructs, no apparent change in neutralizing specificity was apparent in the DMS data for these stabilized and non-stabilized HAs. For all constructs except H5-RC_I_1, it is likely that sufficient antibodies targeting residues adjacent to the RBS were elicited, resulting in the majority of neutralizing activity being directed toward this region, irrespective of HA stabilization.

**Fig. 5.**
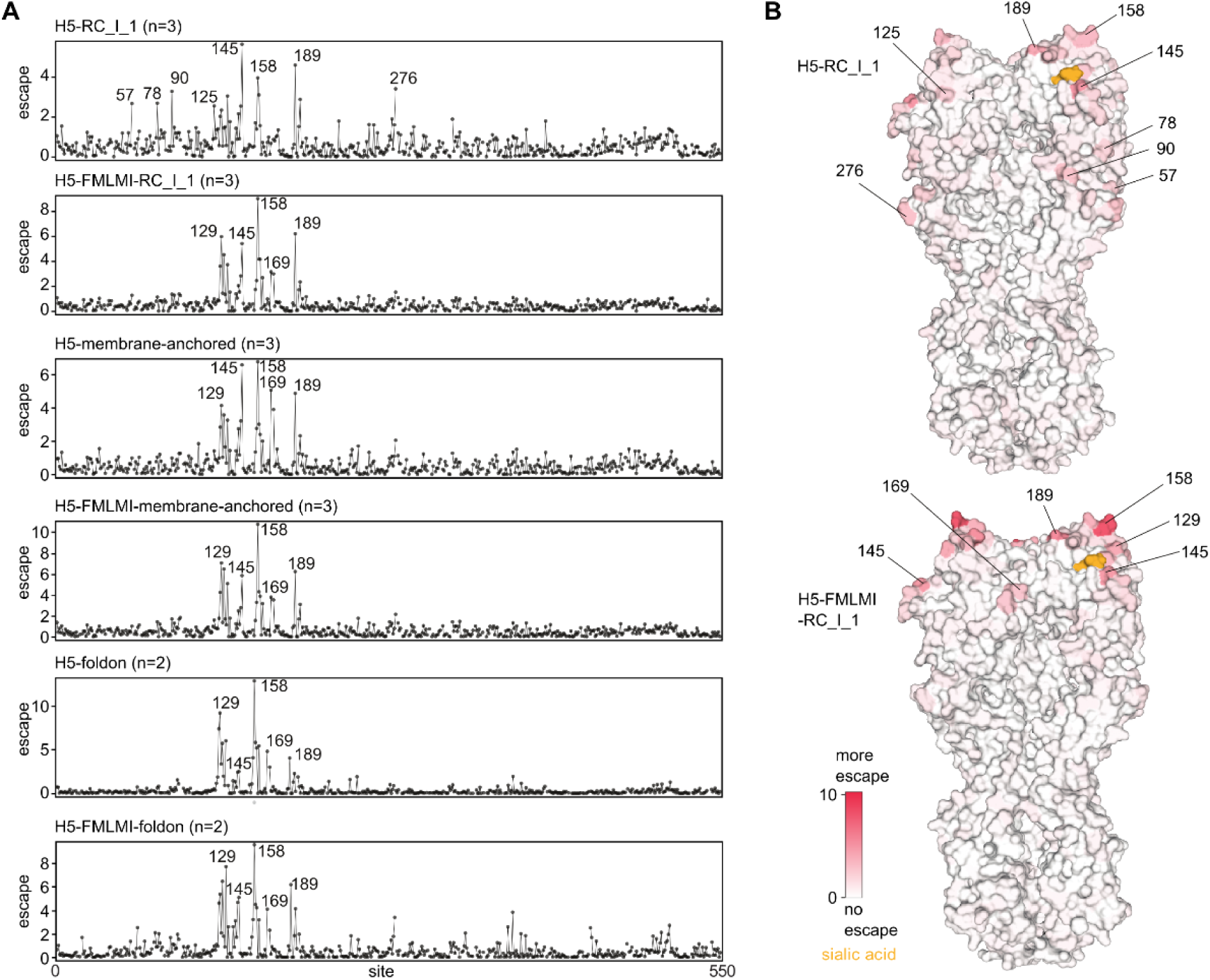
DMS Epitope Mapping of Vaccine-elicited Neutralizing Antibody Responses (A) Escape from serum antibody neutralization for all tolerated amino-acid mutations at each site in HA. Average escape for all animals in each group is shown with the number of sera from each group of animals analyzed indicated in brackets. See https://dms-vep.org/Flu_H5_American-Wigeon_South-Carolina_2021-H5N1_DMS/htmls/Stabilized_HA_sera_escape_faceted.html for individual animal escape measurements and interactive plots. (B) Average serum escape per position for animals from the H5-RC_I_1 and H5-FMLMI-RC_I_1 groups overlaid on HA surface representation (PBD ID: 4KWM). Numbering based on mature H3 HA.

## DISCUSSION

Here we show that stabilizing mutations dramatically enhance thermal stability and improve the quality of vaccine-elicited antibody responses in mice across multiple H5 HA immunogens delivered both as protein and LNP-formulated mRNA. While all soluble HA constructs formed trimeric, prefusion HA0 molecules by nsEM, the stabilizing FMLMI mutations diminished non-neutralizing FluA-20 binding in both the foldon and membrane-anchored groups, suggesting more efficient closed trimer formation in the head domains of these constructs. This resulted in a reduction of internally-directed antibodies elicited by the stabilized immunogens *in vivo*, with a simultaneous boosting of neutralizing RBS-directed responses. We have observed this phenomenon previously, where stabilized, trimeric RBD nanoparticles elicited higher neutralizing responses across several H1 strains than monomeric RBD nanoparticles, and only the latter induced high amounts of internally-directed antibodies (*38*). In addition to epitope (in)accessibility, the considerably improved stability we observed in the FMLMI immunogens may also be a key driver of their enhanced immunogenicity. Previous studies employing series of HIV envelope (*39, 40*) and RSV F (*41*) immunogens showed positive correlations between increasing physical stability and vaccine-elicited neutralizing responses. Moreover, though we did not explore antigen retention studies here, recent work has shown that proteolytic degradation in lymph nodes is a major hurdle for subunit vaccines to overcome in order to reach follicles where adaptive immune responses are generated (*42*). Thus, the correlation between physical stability and enhanced immunogenicity may result from the extension of antigen integrity *in vivo*. Consistent with this, we observed by ns-EMPEM a high amount of site E-binding mAbs across three different H5 HA immunogen formats delivered via mRNA-LNP. The immunodominance of this site in this naïve BALB/c mouse model mirrors the targets of anti-HA B cell hybridomas generated early in the response to H1N1 infection in the same type of mice (*43*). In that study, responses to antigenic sites A and B near the RBS only emerged later during the course of infection. By contrast to infection, where newly produced virions produce a constant supply of new antigen, vaccination in the present study was limited to delivery of new antigen at two discrete time points. This further supports the idea that the stabilizing FMLMI mutations prolong intact antigen availability *in vivo*, so as to generate not only initial responses against the immunodominant site E, but also an increase in potentially later responses against the more neutralizing RBS (*44*).

The stabilizing FMLMI mutations enhanced immunogenicity regardless of construct design and delivery modality, highlighting the utility of incorporating these mutations into any type of HA-based influenza vaccine. This result is particularly timely given the pandemic threat of H5 HPAI viruses currently circulating extremely widely in birds and mammals. Incorporating stabilizing mutations into H5 HA vaccine platforms that can be rapidly produced and scaled up could be an important pandemic preparedness measure (*45*). Although stabilizing mutations were only tested in the context of H5 HA in this study, Milder *et al.* showed substantial stability improvements by incorporating similar mutations across HAs from type A strains (*23*). This further suggests that vaccines targeting any influenza A subtype may be improved by incorporating these universal stabilizing mutations.

More broadly, antigen multimerization in the mRNA-LNP-delivered stabilized nanoparticle and both membrane-anchored formats enhanced immunogenicity compared to soluble HA trimers, a finding that has been observed previously (*17*). This effect is further underscored by the observation that H5-FMLMI-foldon had considerably higher expression than H5-FMLMI-RC_I_1 when produced in HEK293F cells, suggesting that the improved immunogenicity of the H5-FMLMI-RC_I_1 nanoparticle is not simply a result of increased protein expression for mRNA-LNP-delivered immunogens. Furthermore, H5-FMLMI-RC_I_1 elicited significantly higher HAI titers than all other groups when delivered via mRNA-LNP, while HAI titers were similar across all protein-delivered groups. Together, these results highlight the synergistic benefit of combining multivalent antigen display with mRNA-LNP-delivered vaccines. However, the prominent instability of H5-RC_I_1, which was less stable than H5-foldon, suggests that the RC_I_1 nanoparticle scaffold is suboptimal and may constrain the immunogenicity of RC_I_1-based nanoparticle immunogens. Despite this scaffold instability, stabilized H5-FMLMI-RC_I_1 elicited neutralizing responses comparable to those of H5-FMLMI-membrane-anchored. It follows from these data that display of stabilized HA on a better nanoparticle scaffold may further improve immunogenicity. Together, our data underscore that for protein nanoparticle or mRNA-launched nanoparticle vaccines, several features of both the antigen and the scaffold—including conformation, stability, expression, and secretion—should be maximized in order to optimize vaccine efficacy.

## MATERIALS AND METHODS

### Study Design

The main objective of this study was to evaluate the impact of introducing stabilizing mutations in H5 HA on immunogenicity. Multiple HA constructs were tested both with and without stabilizing mutations in a preclinical mouse model. As multimerization is a known driver of immunogenicity, we compared these stabilizing mutations in the context of both soluble trimer and protein nanoparticle immunogens. Additionally, we assessed these immunogens both as protein- and mRNA-LNP-delivered vaccines. By immunizing as protein, we were able to give the same amount of HA across all groups, allowing us to evaluate vaccine constructs independent of expression level. By contrast, delivery with a dose-matched amount of mRNA-LNP allowed us to evaluate these constructs in the rapidly manufacturable and highly scalable mRNA-LNP platform. Since membrane-anchored HA is being widely evaluated as an mRNA-LNP influenza vaccine, we included both stabilized and unstabilized versions of membrane-anchored H5 HA. Two immunizations were administered (weeks 0 and 4) in order to analyze immune responses post-prime (week 2) and post-boost (week 6). ELISA binding, HAI, and microneutralization using both vaccine-matched and vaccine-mismatched H5 strains were performed in order to assess functionality, potency, and breadth of the vaccine-elicited immune responses. ns-EMPEM was used to visualize antibody binding epitopes, while DMS was used to evaluate epitopes targeted by neutralizing antibodies.

All groups consisted of 10 mice to maximize statistical power to assess differences between groups. However, blood from one mouse in the mRNA-LNP-delivered H5-RC_I_1 group was unable to be drawn at week 6, so there are only data from 9 mice in this group at this time point. Animals were randomly assigned to groups and no blinding was implemented.

While all HA numbering in this paper is based on H3 HA numbering, conversions to other HA numbering schemes can be found here: https://dms-vep.org/Flu_H5_American-Wigeon_South-Carolina_2021-H5N1_DMS/numbering.html.

### Gene Expression and Protein Purification

All HA constructs used in this study were codon-optimized for human cell expression and made in the CMV/R vector (*46*) by Genscript with a C-terminal hexahistidine affinity tag. PEI MAX was used for transient transfection of HEK293F cells. After four days, mammalian cell supernatants were clarified via centrifugation and filtration. HA protein immunogens and HA foldons were all purified using IMAC. 1 mL of Ni^2+^-sepharose Excel was added per 100 mL clarified supernatant along with 5 mL of 1 M Tris, pH 8.0 and 7 mL of 5 M NaCl and left to batch bind while shaking at room temperature for 30 min. Resin was then collected in a gravity column, washed with 5 column volumes of 50 mM Tris, pH 8.0, 500 mM NaCl, 20 mM imidazole, and protein was eluted using 50 mM Tris, pH 8.0, 500 mM NaCl, 300 mM imidazole. Further purification was done using SEC on a Superose 6 Increase 10/300 gel filtration column equilibrated in 25 mM Tris, pH 8.0, 150 mM NaCl, 5% glycerol. HA-ferritin nanoparticles used in HAI assays were purified using lectin affinity chromatography, followed by SEC on a Superose 6 Increase 10/300 gel filtration column (*47*).

Following purification, nanoparticle quality and concentration was first measured by UV-vis spectroscopy. Nanoparticle polydispersity and purity post freeze-thaw was then assessed using SDS-PAGE, DLS, and nsEM. Finally, endotoxin levels were measured using the LAL assay, with all immunogens used in animal studies containing less than 100 EU/mg in the final dose. Final immunogens were flash-frozen using liquid nitrogen and stored at −80°C.

### Flow Cytometry

HEK293F cells were transiently transfected with HA-membrane-anchored constructs. Following a wash with PBS, Zombie Aqua live/dead viability dye (BioLegend) and 1 μM biotinylated FluA-20 was added to cells and left to incubate for 15 min. Cells were again washed and subsequently stained with Streptavidin, R-Phycoerythrin Conjugate (SAPE; Thermo Fisher) and 200 nM AF647-conjugated CR9114 for another 15 min. Following a final wash, flow cytometry was performed using an Attune flow cytometer (Thermo Fisher). The Live/Singlets populations were used in final binding plots.

### NanoDSF

All proteins were formulated at 0.5 mg/mL in 25 mM Tris, pH 8.0, 150 mM NaCl, 5% glycerol and then mixed at 9 volumes to 1 volume of 200× concentrate SYPRO orange (Thermo Fisher) diluted in the same buffer.

NanoDSF to determine melting temperatures was carried out on an UNcle (UNchained Labs) by measuring the integration of fluorescence emission spectrum during a thermal ramp from 25°C to 95°C, with a 1°C increase in temperature per minute.

### Bio-layer Interferometry

BLI was carried out using an Octet Red 96 system, at 25°C with 1000 rpm shaking. Anti-HA antibodies were diluted in kinetics buffer (PBS with 0.5% serum bovine albumin and 0.01% Tween) to a final concentration of 10 μg/mL before loading onto protein A biosensors (Sartorius) for 200 sec. HA proteins were diluted to 500 nM in kinetics buffer and their association was measured for 200 sec, followed by dissociation for 200 sec in kinetics buffer alone.

### Negative Stain Electron Microscopy

3.5 µL of 70 µg/mL nanoparticles were applied to glow-discharged 400-mesh carbon-coated grids (Electron Microscopy Sciences) and stained with 2% (wt/vol) uranyl formate. Data were collected using EPU 2.0 on a 120 kV Talos L120C transmission electron microscope (Thermo Scientific) with a BM-Ceta camera. CryoSPARC (*48*) was used for CTF correction, particle picking and extraction, and 2D classification.

### Dynamic Light Scattering

DLS was carried out on an UNcle (UNchained Labs) at 25°C. 10 acquisitions of 5 sec each were acquired for each spectrum. Protein concentration (ranging from 0.1-1 mg/mL) and buffer conditions were accounted for in the software.

### Mass Photometry

MP was carried out in a TwoMP (Refeyn) Mass photometer. A 12-well gasket was placed on each slide. 10 μL of buffer was added to one well of this gasket and the camera was brought into focus after orienting the laser to the center of the sample well. Next, 2 μL of H5-FMLMI-RC_I_1 at 120 nM nanoparticle was added to this droplet and 1 min videos were collected with a large field of view in AcquireMP. Ratiometric contrast values for individual particles were measured and processed into mass distributions with DiscoverMP. A sample of 20 nM Beta-amylase—consisting of monomers (56 kDa), dimers (112 kDa) and tetramers (224 kDa) in equilibrium—was used to arrive at a mass calibration, thereby allowing contrast values to be converted into mass values. Distributions and Gaussian fit were plotted using the seaborn Python library (*49*).

### Cryo-EM sample preparation, data collection and processing

Cryo-EM samples were prepared on glow-discharged QUANTIFOIL 1.2/1.3 holey-carbon grids (Electron Microscopy Sciences). Protein sample at 1 mg/ml was applied to the grid, incubated for 30 seconds, and then blotted using Whatman filter paper; this process was performed twice, followed by a third application of sample and final blotting and plunge freezing in liquid-ethane using a Vitrobot (FEI). Data were acquired on a Titan Krios operating at 300 kV equipped with a K3 Summit Direct Detector and a Quantum GIF energy filter (Gatan) operating in zero-loss mode with a 20-eV slit width. Movies were collected at both 0° and 30°, in counting mode, collecting 100 frames with a total dose of 60 e−/Å−2, with a defocus range of −0.5 to −2 µm. Automated data collection was carried out using SerialEM(*50*) at a nominal magnification of 105,000× with a pixel size of 0.843 Å. All data processing was carried out using cryoSPARC(*48*). Movie frame alignment and dose-weighting summation were performed using Patch Motion, and CTF estimation was done using patch CTF. Particles were first picked using Blob picker and these were used to generate initial 2D averages. Two more rounds of 2D classification were performed to select all well-aligned, single HA views, followed by a final selection of only HA side views. Top views of HA were excluded due to their preferred orientation and thus potential damage at the air-water interface. An initial model was generated by ab-initio reconstruction with C1 symmetry, followed by an initial homogeneous 3D refinement with C3 symmetry. A solvent mask around the HA trimer was then used to exclude noise in a 3D classification without alignment, and in all subsequent steps. The final homogeneous refinement of the 3D class with the most particles resulted in the best-resolved map at 3.2 Å and was used to build an atomic model.

### Model building and refinement

An AlphaFold3(*51*) model was docked into the cryo-EM map as a rigid body using Chimera(*52*). A single N-acetylglucosamine at each non-clashing glycan sequon was added using GlycoShape(*53*). An initial refinement of one HA protomer into the cryo-EM density was performed with ISOLDE(*54*). C3 symmetric copies of this fit protomer were then generated and this HA trimer model was relaxed into the cryo-EM density again in ISOLDE. A final real-space refinement with grid searching, Ramachandran and rotamer restraints all turned off, and ‘use starting model as reference’ turned on, was then performed in Phenix(*55*). Flexible loops comprising residues XX-XX and XX-XX, as well as the foldon domain, were not resolved in the cryo-EM density so these loops were not modeled.

### mRNA Design and Production

The H5 HA-encoding mRNAs were produced as described(*21*). Briefly, the H5 HA sequences were codon-optimized, synthesized, and cloned into an mRNA production plasmid. The mRNA was transcribed to contain a 101 nucleotide-long poly(A) tail. N1-methylpseudouridine-5′-triphosphate (m1Ψ-5′-triphosphate) (TriLink #N-1081) instead of uridine-5′-triphosphate (UTP) was used to generate modified nucleoside-containing mRNA. Capping of the *in vitro* transcribed mRNAs was performed co-transcriptionally using the trinucleotide cap1 analog, CleanCap (TriLink #N-7413). mRNA was purified by cellulose (Sigma-Aldrich #11363-250G) purification. The mRNAs were analyzed by agarose gel electrophoresis and were stored frozen at −20°C.

### LNP Encapsulation of mRNA

mRNA-LNP, with similar composition was produced as described previously (*21*), which contains cationic ionizable lipid and PEGylated lipid proprietary to Acuitas Therapeutics. The lipids and formulation are described in patent application WO 2017/004143A1. In brief, ionizable lipid/distearoylphosphatidylcholine (DSPC)/cholesterol/PEGylated lipid were dissolved in ethanol and rapidly mixed with mRNA diluted in an aqueous citrate pH 4 buffer. Formulations were then dialyzed overnight with PBS, concentrated to target 1 mg/mL with cryoprotectant, filtered, vialed and then frozen at −80°C. mRNA-LNP was characterized at Acuitas Therapeutics for their size, polydispersity using a Malvern Zetasizer (Malvern Panalytical), encapsulation efficiency using the standard Ribogreen assay (Thermo Fisher Scientific), and mRNA content by UPLC analysis (Waters).

### Ethical Statement

Animal experiments were conducted in accordance with the University of Washington’s Institutional Animal Care and Use Committee (IACUC protocol 4470-01). The University of Washington’s animal use program is accredited by the Association for Assessment and Accreditation of Laboratory Animal Care (Reference Assurance: #000523).

### Immunization

Female BALB/c mice were purchased from Envigo (order code 047) at 7 weeks of age and were housed in a specific pathogen-free facility within the Department of Comparative Medicine at the University of Washington, Seattle. Animals had access to standard rodent chow and water *ad libitum* and were maintained in an ABSL1 housing room with environmental parameters set to 68-79°F, 30-70% humidity, and a dark/light cycle of 10 hours of dark and 14 hours of light. At 8 weeks of age, mice were immunized subcutaneously in the inguinal region with 21.8 pmol protein in a 50% (v/v) mixture of AddaVax adjuvant (InvivoGen, San Diego, CA) or intramuscularly with 1 μg mRNA-LNP formulated in 100 μL PBS (50 μL in each quadricep) at weeks 0 and 4. For mice receiving IM injections, preemptive 0.05mg/kg Buprenorphine HCl (NDC 42023-179-05) was administered subcutaneously to prevent pain. For sera, animals were bled using the submental route at week 0 and 2 weeks post-immunization. Whole blood was collected in serum separator tubes (BD #365967) and rested for 30 min at room temperature for coagulation. Tubes were then centrifuged for 10 min at 2,000 *g* and serum was collected and stored at −80°C until use.

### ELISA

H5-I53_dn5B trimers were added to 96-well half-area high binding plates (Corning) at 2.0 μg/mL with 25 μL per well and incubated overnight at 4°C. Blocking buffer composed of Tris Buffered Saline Tween (TBST: 25 mM Tris pH 8.0, 150 mM NaCl, 0.05% (v/v) Tween20) with 5% Nonfat milk was then added at 100 μL per well and incubated for 1 hour. Next plates were washed, with all washing steps consisting of 3× washing with TBST using a robotic plate washer (Biotek). 5-fold serial dilutions of serum starting at 1:100 were made in blocking buffer, added to plates at 25 μL per well, and incubated for 1 hour. Plates were washed again before addition of 25 μL per well of anti-mouse HRP-conjugated secondary antibody (CellSignaling Technology) diluted 1:2,000 in blocking buffer and incubated for 30 minutes. All incubations were carried out with shaking at room temperature. Plates were washed a final time, and then 50 μL per well of TMB (3,3′,5′,5-tetramethylbenzidine, SeraCare) was added for 5 minutes, followed by quenching with 50 μL per well of 1 N HCl. Reading at 450 nm absorbance was done on an Epoch plate reader (BioTek).

### HAI

Serum was inactivated using receptor-destroying enzyme (RDE) II (Seiken) in PBS at a 3:1 ratio of RDE to serum for 16 hours at 37°C, followed by 40 minutes at 56°C. Inactivated serum was serially diluted 2-fold in PBS in V-bottom plates at 25 μL per well. 25 μL HA-ferritin nanoparticles at 4 hemagglutinating units were then added to all wells and incubated at room temperature for 30 min (*30*). Lastly, 50 μL of 10-fold diluted turkey or horse red blood cells (Lampire) in PBS was added to each well. Hemagglutination was left to proceed for at least 1 hour before recording HAI titer.

### Reporter-based Microneutralization Assay

Influenza A reporter viruses were made as previously described (*56*). Briefly, H5N1 viruses were made with a modified PB1 segment expressing the TdKatushka reporter gene (R3ΔPB1), rescued, and propagated in MDCK-SIAT-PB1 cells in the presence of TPCK-treated trypsin (1 μg/mL, Sigma) at 37°C. Virus stocks were stored at −80°C and were titrated before use in the assay. Sera was treated with receptor destroying enzyme (RDE II; Denka Seiken) and heat-inactivated before use in neutralization assays. 384 well plates (Greiner) were pre-seeded with 1.0 × 10^5^ MDCK-SIAT1-PB1 cells and incubated overnight. Immune sera were serially diluted and incubated for 1 h at 37°C with pre-titrated virus (A/Texas/37/2024 and A/Indonesia/175H/2005).

Serum-virus mixtures were then transferred in quadruplicate onto the pre-seeded 384 well plates and incubated at 37°C for 18-26 hours. The number of fluorescent cells in each well was counted automatically using a Celigo image cytometer (Nexcelom Biosciences). IC_50_ values, defined as the serum dilution or antibody concentration that gives 50% reduction in virus-infected cells, were calculated from neutralization curves using a four-parameter nonlinear regression model.

### ns-EMPEM

50 μL serum from each of the 10 mice per group was pooled and then mixed with an equal volume of Acetate buffer, pH 5.0, and incubated overnight with 0.5 ml packed Protein G agarose resin (Thermo Fisher). The flowthrough was removed using gravity purification and resin was washed with 20 CVs PBS. IgGs were eluted by incubating resin for 20 min with 1 CV of 0.1 M glycine, pH 2.5, repeated twice. Elutions were neutralized with 1 M Tris, pH 8.0 to a final concentration of 50 mM. IgGs were then buffer exchanged into PBS and concentrated to 200 μL for digestion into Fabs. Fab digestion was carried out by adding 200 μL freshly made 2× digestion buffer (40 mM NaPO_4_ pH 6.5, 20 mM EDTA, 40 mM Cysteine) and 300 μL papain resin in 300 μL 1× digestion buffer, and incubated with shaking at 37°C for 16 hours. Papain digestion reaction was centrifuged and supernatant containing Fabs was collected and filtered. Papain was then washed with 1 CV 20 mM Tris, pH 8.0 and this supernatant was added to the first. Fabs were then concentrated to 50 μL and mixed at a 50-fold molar excess to H5-I53_dn5B trimer, and incubated for 16-20 hours at room temperature with gentle rocking. Immune complexes were purified by SEC on a Superdex 200 Increase 10/300 GL column and used in EM. nsEM data were collected using EPU 2.0 on a 120 kV Talos L120C transmission electron microscope (Thermo Scientific) with a BM-Ceta camera. Data processing was done in CryoSPARC (*48*), starting with CTF correction, particle picking and extraction. Three rounds of 2D classification were done, keeping only classes that had either HA monomer or trimer with bound Fabs. 3D models for these immune complexes were then generated using ab initio 3D reconstruction and heterogeneous refinement without imposing any symmetry.

Classes that had clear, trimeric HA density and were representative of the diversity of Fab binding in each sample of polyclonal serum without redundancy were then separately subjected to a 3D refinement.

### Deep mutational scanning

H5 HA deep mutational scanning libraries were produced in A/American Wigeon/South Carolina/USDA-000345-001/2021 strain background as described previously (*11*). A/American Wigeon/South Carolina/USDA-000345-001/2021 strain differs from A/Texas/37/2024 strain by 2 mutations (L122Q and T199I, H3 numbering) in the HA gene. To measure how HA mutations affect serum neutralization, deep mutational scanning libraries were incubated for 45 min at 37°C with serum dilutions that neutralize between 80-99% of the library viruses. After incubation with sera, library mixtures were used to infect 293T cells and 15 h post infection viral genomes were recovered and prepared for Illumina sequencing. One biological replicate was performed for each sera. Before use all sera was treated with receptor destroying enzyme for 2 h at 37°C and subsequently heat inactivated at 56°C for 30 min. A biophysical model implemented in *dms-vep-pipeline-3* was used to calculate escape caused by each mutation in the libraries (*57*). Interactive plots for serum escape data are available at https://dms-vep.org/Flu_H5_American-Wigeon_South-Carolina_2021-H5N1_DMS/htmls/Stabilized_HA_sera_escape_faceted.html. Analysis pipeline used to model serum escape is available on GitHub https://github.com/dms-vep/Flu_H5_American-Wigeon_South-Carolina_2021-H5N1_DMS. The plots in Fig. 5 show the total escape caused by all tolerated amino acids at each site, averaging over mice in each group.

### Statistical analysis

Multi-group comparisons were performed using the Brown-Forsythe one-way ANOVA test and Dunnett’s T3 post hoc analysis in Prism 9 (GraphPad). Differences were considered significant when p values were less than 0.05.

## Acknowledgements

We thank S. Ols for assistance with flow cytometry data acquisition.

## Author Contributions

Conceptualization: AD and NPK

Investigation: AD, BD, RG, EML, JM, EG, MV, HM, and RJ

Funding acquisition: MK, JDB, and NPK

Supervision: MK, JDB, and NPK Writing – original draft: AD and BD

Writing – reviewing and editing: AD, BD, MK, JDB, and NPK

## Data and materials availability

All data needed to evaluate the conclusions in the paper are present in the paper and/or the Supplementary Materials. nsEM maps have been deposited at the Electron Microscopy Data Bank (EMDB) with accession codes EMD-71459 (Fab 3 reconstruction of polyclonal serum from mouse immunized with H5 TX24-FMLMI-RC_I_1 in complex with TX24-I53_dn5B), EMD-71460 (Fab 2 reconstruction of polyclonal serum from mouse immunized with H5 TX24-FMLMI-RC_I_1 in complex with TX24-I53_dn5B), EMD-71461 (Fab 1 reconstruction of polyclonal serum from mouse immunized with H5 TX24-FMLMI-RC_I_1 in complex with TX24-I53_dn5B), EMD-71462 (Fab reconstruction of polyclonal serum from mouse immunized with H5 TX24-foldon in complex with TX24-I53_dn5B), EMD-71463 (Fab 1 reconstruction of polyclonal serum from mouse immunized with H5 TX24-FMLMI-foldon in complex with TX24-I53_dn5B), EMD-71464 (Fab 2 reconstruction of polyclonal serum from mouse immunized with H5 TX24-FMLMI-foldon in complex with TX24-I53_dn5B), EMD-71465 (Fab 3 reconstruction of polyclonal serum from mouse immunized with H5 TX24-FMLMI-foldon in complex with TX24-I53_dn5B), EMD-71466 (Fab 1 reconstruction of polyclonal serum from mouse immunized with H5 TX24-membrane-anchored in complex with TX24-I53_dn5B), EMD-71467 (Fab 2 reconstruction of polyclonal serum from mouse immunized with H5 TX24-membrane-anchored in complex with TX24-I53_dn5B), EMD-71468 (Fab 3 reconstruction of polyclonal serum from mouse immunized with H5 TX24-membrane-anchored in complex with TX24-I53_dn5B), EMD-71469 (Fab 4 reconstruction of polyclonal serum from mouse immunized with H5 TX24-membrane-anchored in complex with TX24-I53_dn5B), EMD-71470 (Fab 1 reconstruction of polyclonal serum from mouse immunized with H5 TX24-FMLMI-membrane-anchored in complex with TX24-I53_dn5B), EMD-71471 (Fab 2 reconstruction of polyclonal serum from mouse immunized with H5 TX24-FMLMI-membrane-anchored in complex with TX24-I53_dn5B), EMD-71472 (Fab 3 reconstruction of polyclonal serum from mouse immunized with H5 TX24-FMLMI-membrane-anchored in complex with TX24-I53_dn5B), EMD-71473 (Fab 4 reconstruction of polyclonal serum from mouse immunized with H5 TX24-FMLMI-membrane-anchored in complex with TX24-I53_dn5B). The cryo-EM map and model have been deposited at the EMDB with accession code EMD-71796 and at the RCSB Protein Data Bank (PDB) with accession code 9PR3.

---

**Fig. S1.**
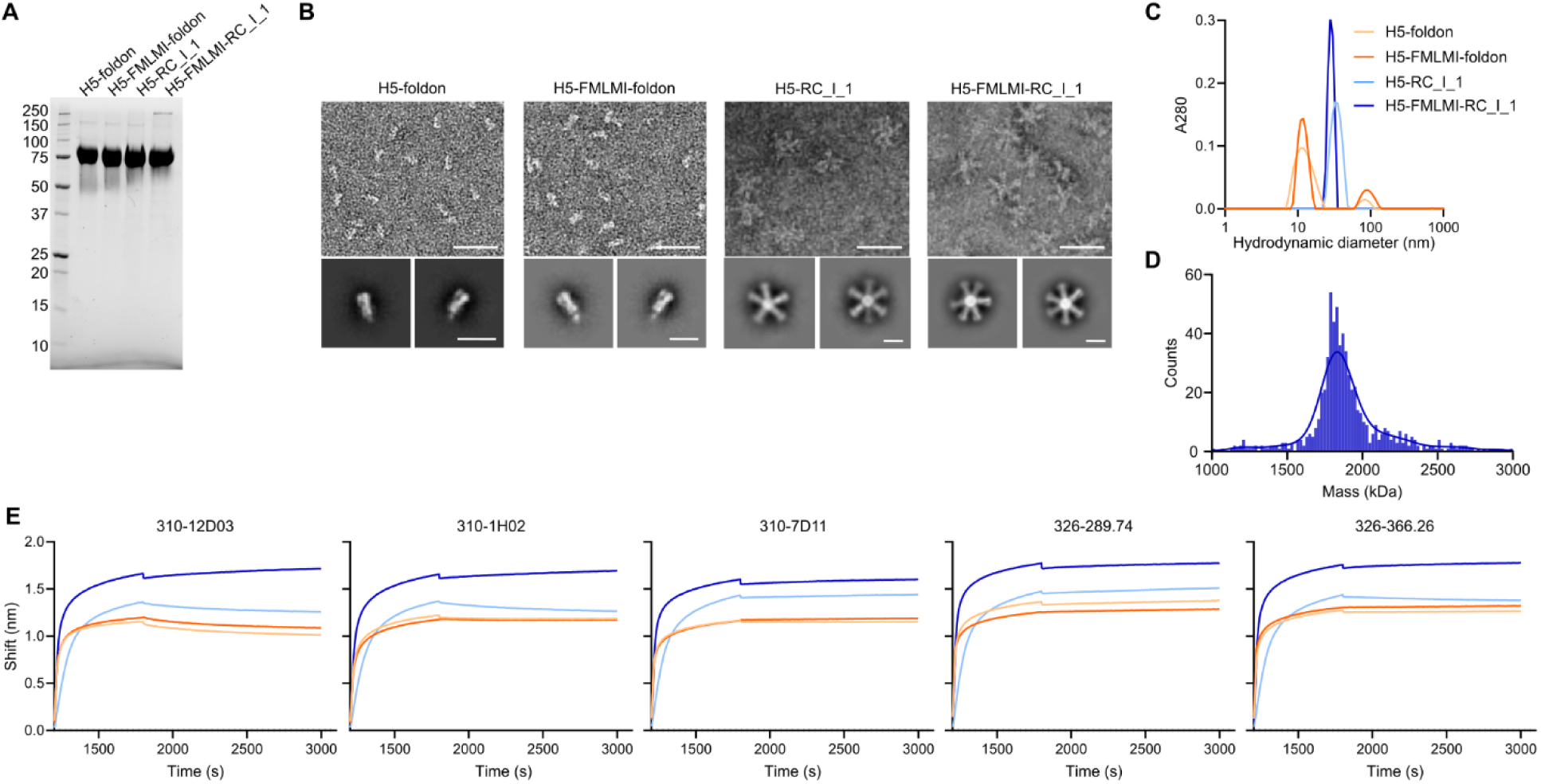
Biophysical characterization of H5-foldon and nanoparticle constructs. (A-C) A. Reducing SDS-PAGE, B. nsEM micrographs and 2D averages, and C. DLS of H5 HA constructs. Micrograph scale bars = 50 nm, class average scale bars = 150 nm. (A) MP histogram and Gaussian fit of H5-FMLMI-RC_I_1. (B) BLI using anti-RBS and anti-vestigial esterase domain mAbs. Same legend as panel C.

**Fig. S2.**
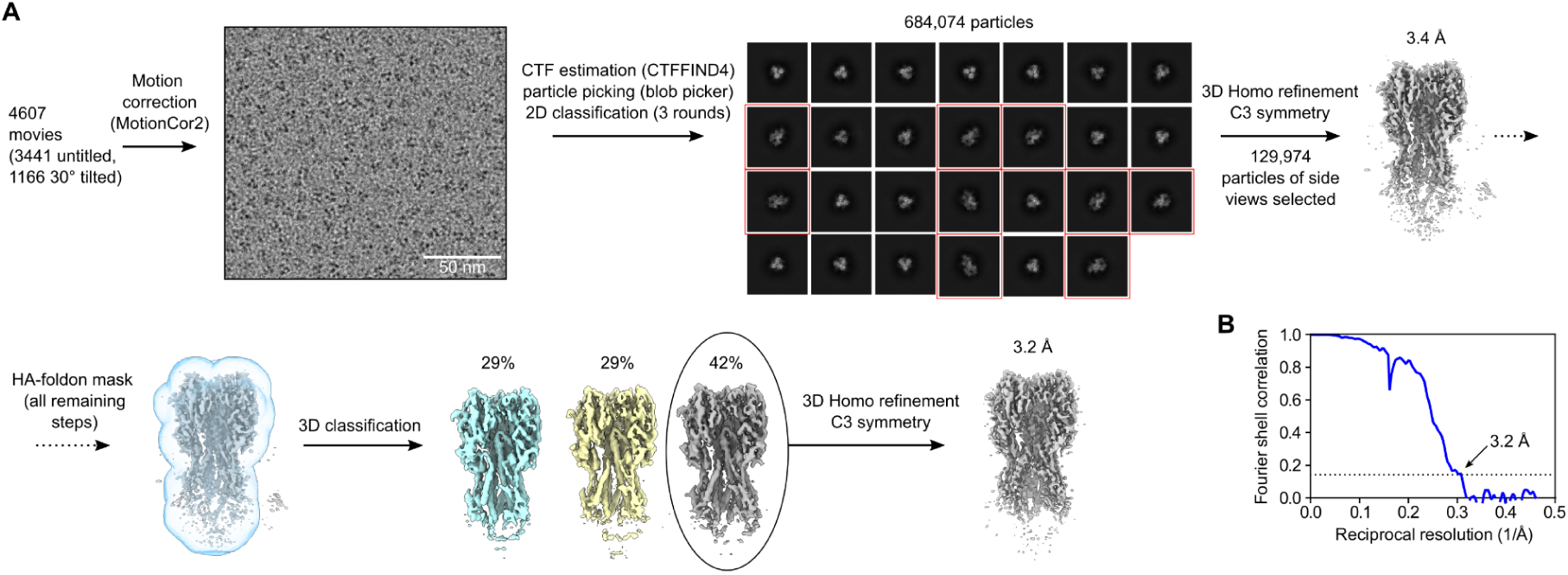
Cryo-EM processing workflow for H5-FMLMI-foldon. (A) H5-FMLMI-foldon reconstruction cryo-EM processing diagram. Red boxes around 2D averages show particles of HA-foldon side views selected for further processing. (B) Gold-standard FSC curve for H5-FMLMI-foldon reconstruction.

**Fig. S3.**
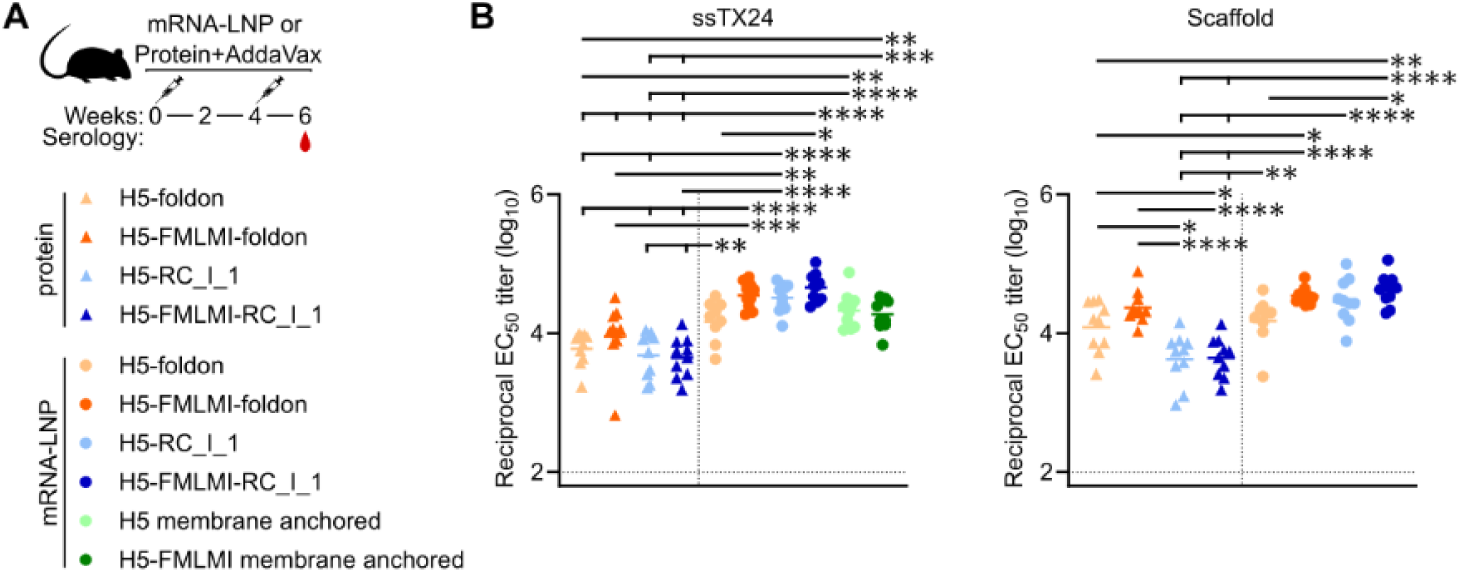
Vaccine-elicited Antibody Responses in Mice Immunized with H5 protein- and mRNA-LNP-delivered constructs (A) H5 mouse immunization schedule and groups. (B) ELISA binding titers in immune sera at week 6 against a stem-only TX24 antigen (left) or RC_I_1 nanoparticle lacking displayed antigen (right). Each symbol represents an individual animal, and the geometric mean of each group is indicated by the bar (N = 5 rabbits/group). Statistical significance was determined using one-way ANOVA with Tukey’s multiple comparisons test; *, p < 0.05; **, p < 0.01; ***, p < 0.001; ****, p < 0.0001.

**Table S1.**
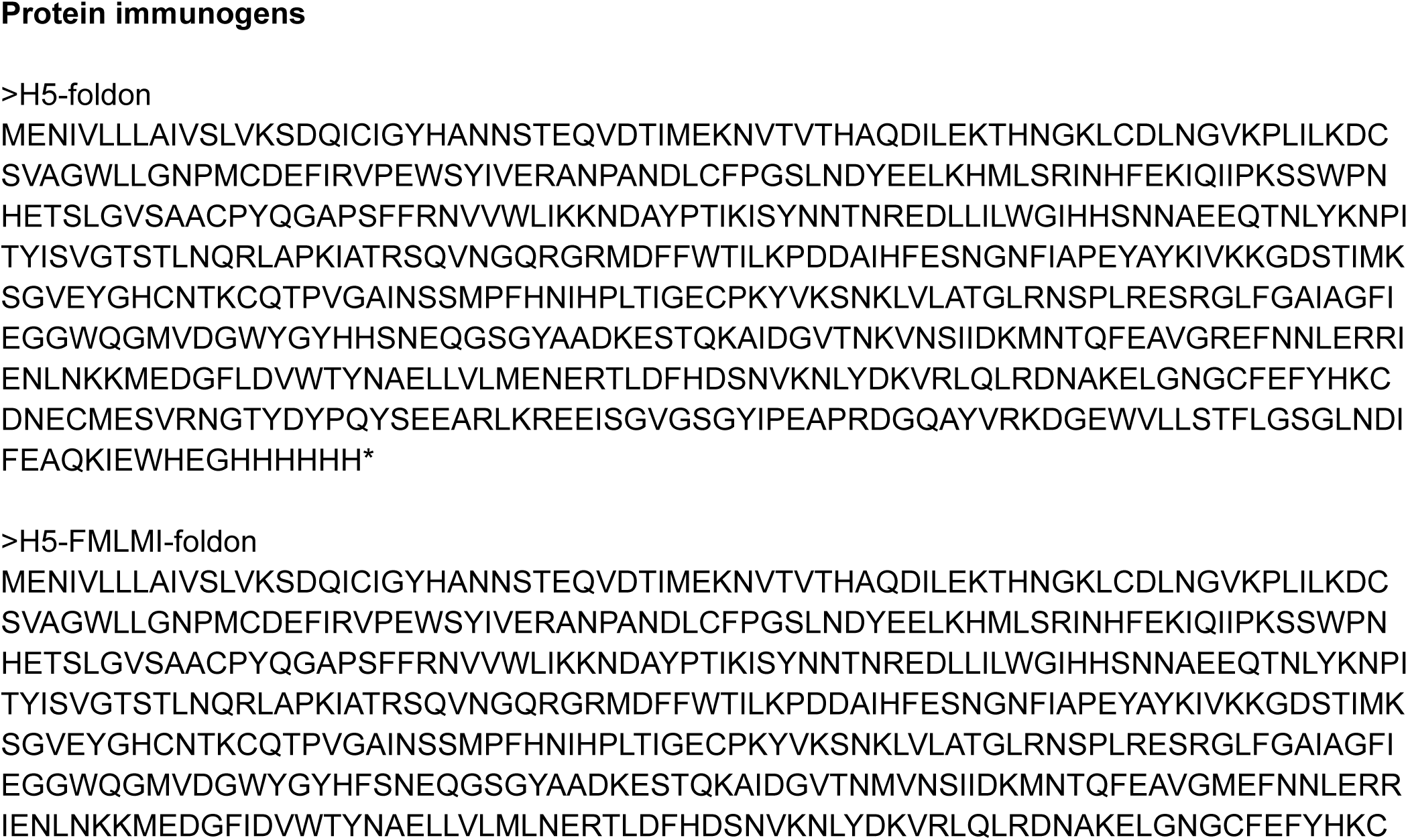
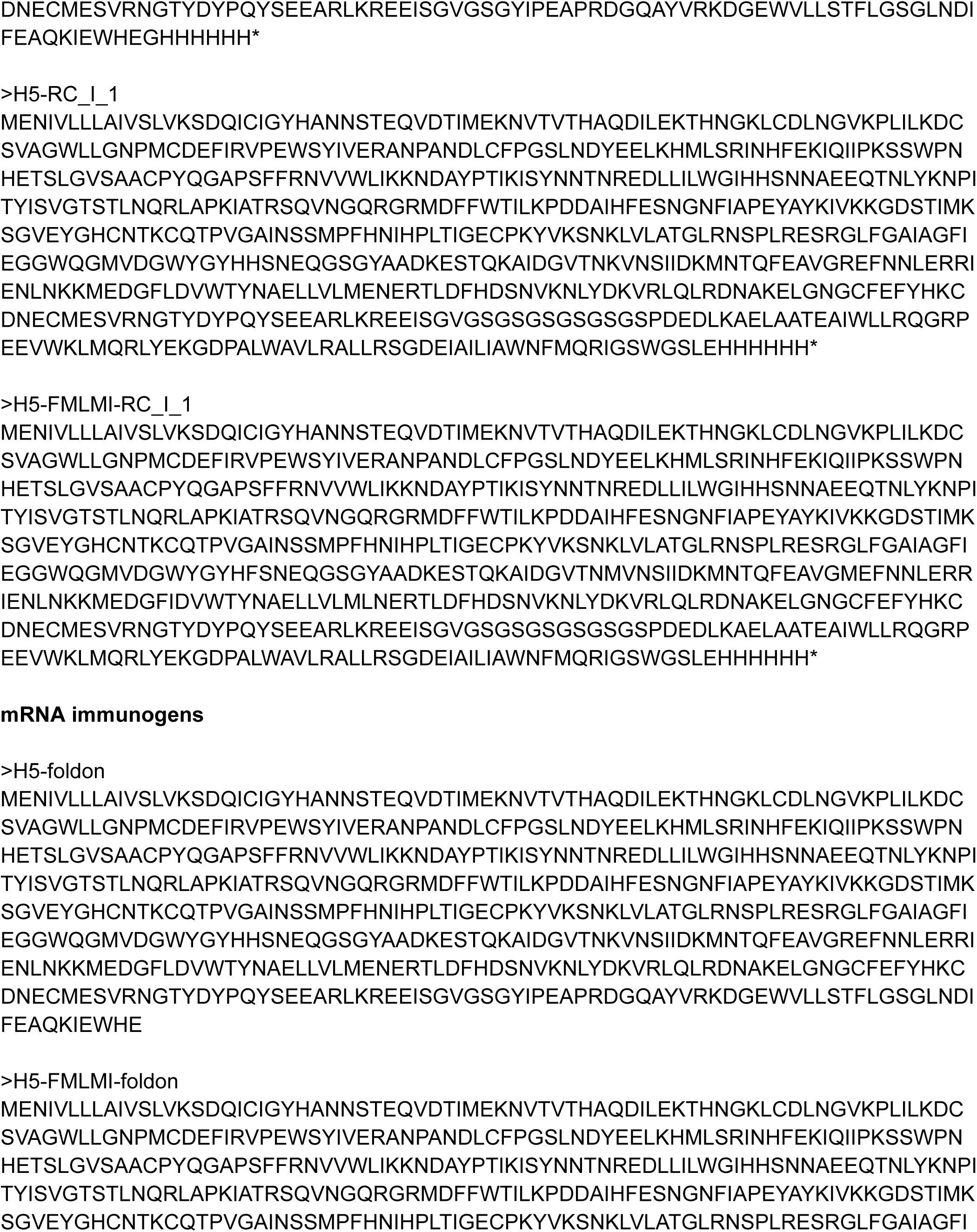
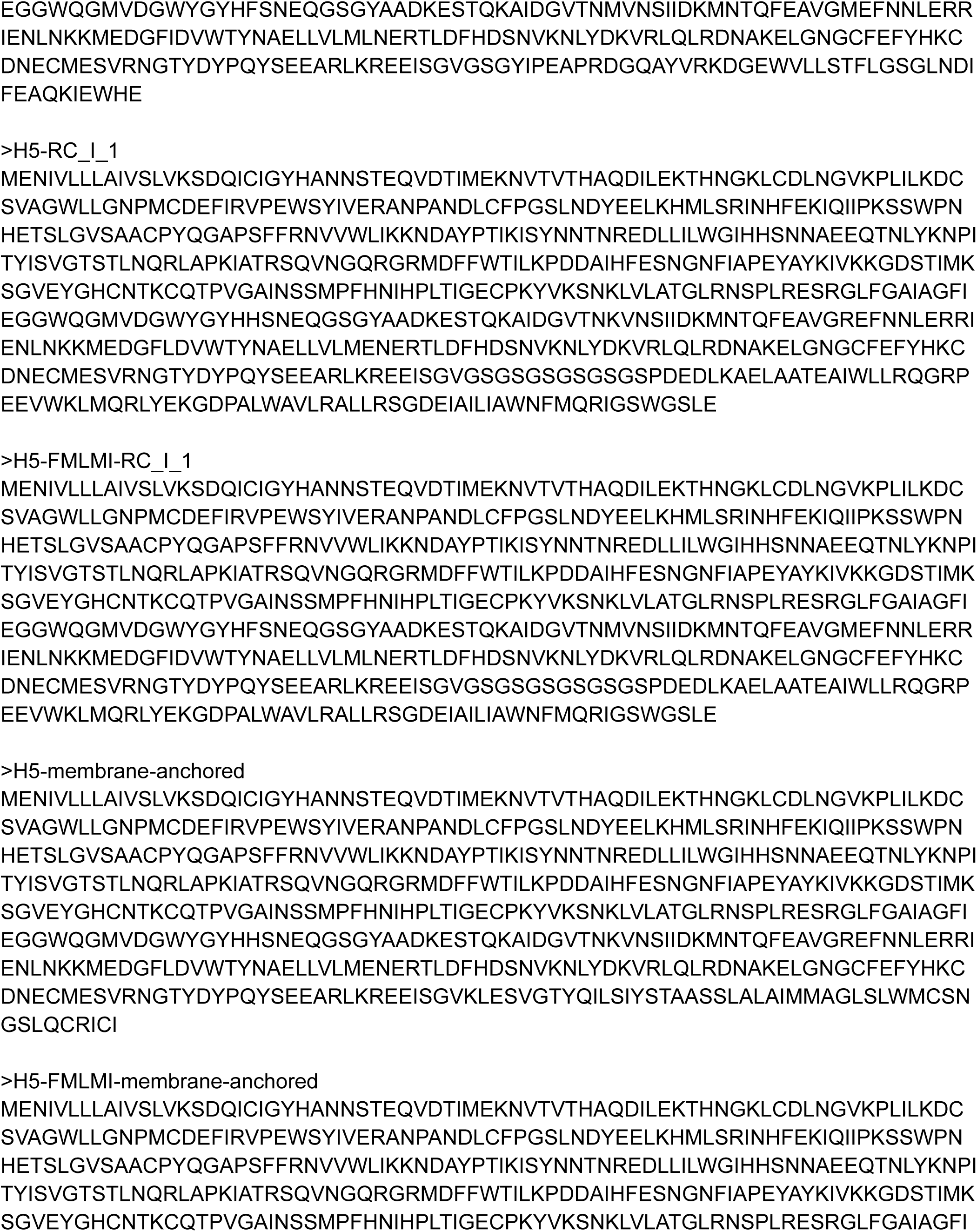
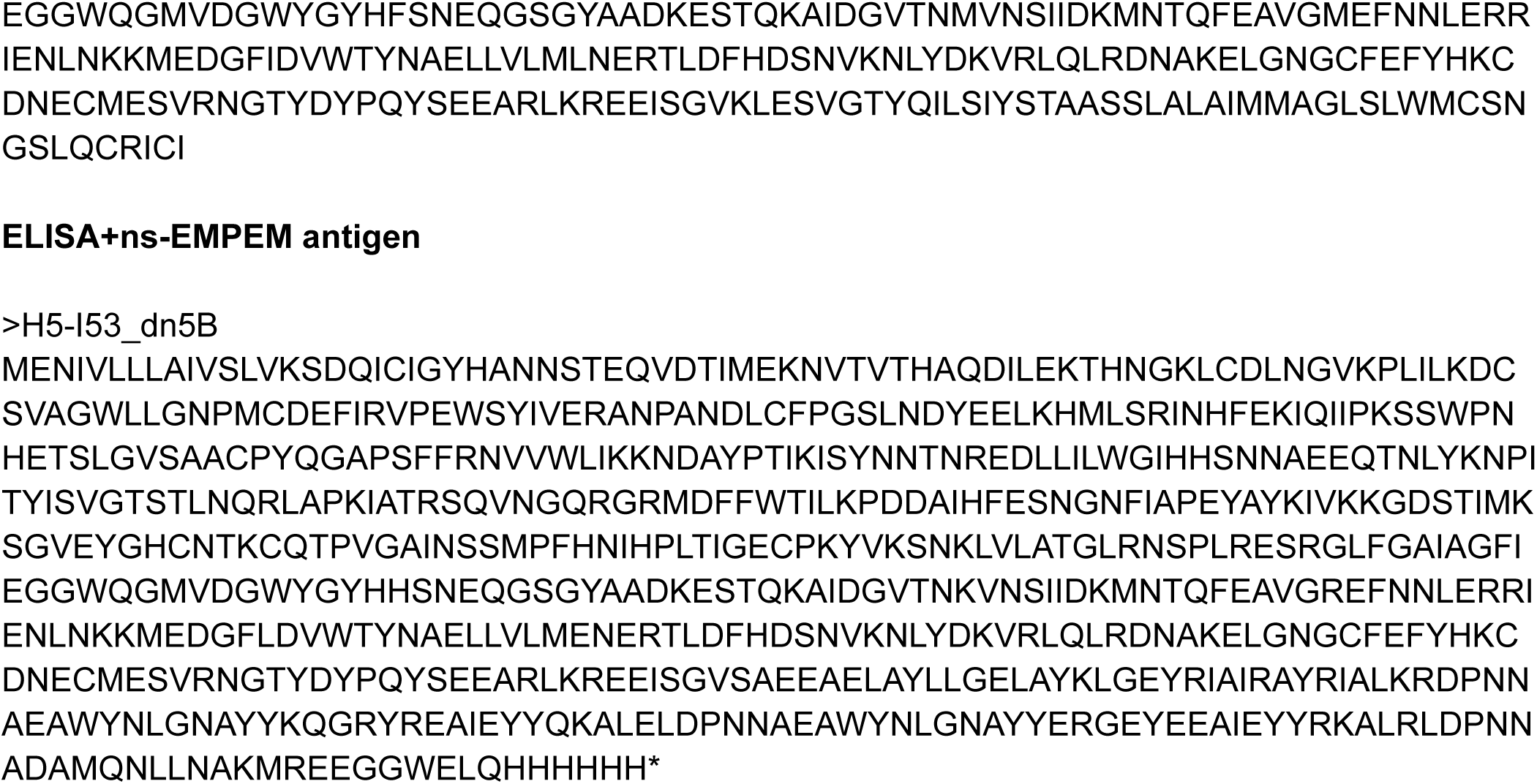
Amino acid sequences of proteins used in this study. Protein immunogens.

**Table S2.** CryoEM data collection and refinement statistics.

|  |  |
| --- | --- |
| H5-FMLMI-foldon<br>PDB 9PR3<br>EMD 71796 |  |
| Data collection and processing |  |
| Magnification (×) | 105,000 |
| Voltage (kV) | 300 |
| Electron exposure (e<sup>-</sup>/Å<sup>2</sup>) | 60 |
| Defocus range (µm) | -0.5 - -2 |
| Pixel size (Å) | 0.843 |
| Symmetry imposed | C3 |
| Final particle images (no.) | 54,035 |
| Map resolution (Å) | 3.2 |
| FSC threshold | 0.143 |
| Map sharpening *B* factor (Å<sup>2</sup>) | -84 |
| Validation |  |
| MolProbity score | 1.96 |
| Clashscore | 8.44 |
| Poor rotamers (%) | 0.5 |
| Ramachandran plot |  |
| Favored (%) | 91.58 |
| Allowed (%) | 8.32 |
| Disallowed (%) | 0.1 |

